# Serine activates defense mechanisms via glutamate receptor-like channel 3.4 and calcium-dependent protein kinase 5 signaling in plants

**DOI:** 10.1101/2025.04.04.647287

**Authors:** Kyounghee Lee, Amanda Navodani, Janint Camacho, Heejin Yoo

## Abstract

Amino acids are fundamental building blocks of proteins, as well as essential metabolites for plant growth and development. Beyond their metabolic roles, amino acids have been increasingly recognized as regulators of plant immunity. However, the molecular mechanisms by which amino acids modulate defense responses remain unclear. In this study, we identify a novel function for serine as a defense-priming signal. Serine pretreatment enhances immune gene expression and primes plants for a more rapid and robust response to pathogen infection, resulting in a significant delay in programmed cell death (PCD) during effector-triggered immunity (ETI). We further demonstrate that serine-induced defense activation is mediated by glutamate receptor-like calcium channel 3.4 (GLR3.4), which subsequently activates calcium-dependent protein kinase 5 (CPK5). CPK5 in turn phosphorylates WRKY transcription factors, including WRKY33, to induce key immune genes. Our findings uncover a previously unrecognized serine-initiated signaling pathway that primes plant defense, highlighting both its mechanistic novelty and its potential application in sustainable agriculture. Given that serine is a naturally occurring metabolite, its exogenous application—or the use of structurally similar analogs—may offer a cost-effective, environmentally friendly, and non-genetic strategy to enhance disease resistance in crops.

## Introduction

Plants, unlike animals, lack specialized immune cells and must reprogram their cellular systems to mount an effective defense against pathogens (Spoel and Dong, 2012). This reprogramming involves dynamic modifications at the transcriptional, translational, and metabolic levels (He et al., 2022; Monson et al., 2022). A well-established aspect of this response is the metabolic shift from growth-related processes toward defense activation, which leads to energy trade-offs that can suppress growth (Rojas et al., 2014). This shift poses a challenge for genetic engineering approaches that aim to enhance disease resistance while maintaining plant productivity (Shah et al., 1999; Kirik et al., 2001).

Recent evidence suggests that primary metabolites, beyond serving as energy sources, actively regulate plant immunity by acting as defense elicitors, highlighting the intricate crosstalk between metabolism and immune responses (Rojas et al., 2014). Among these metabolites, amino acids have gained increasing attention as potential immune modulators (Fabro et al., 2004; Ward et al., 2010; Chen et al., 2011; Yoo et al., 2020). Several studies have demonstrated that amino acid levels fluctuate dynamically during pathogen infection. For instance, proline accumulates in Arabidopsis leaves following infection with avirulent *Pseudomonas syringae* strains, suggesting a role in plant defense (Fabro et al., 2004). Similarly, valine, leucine, isoleucine, and threonine accumulate upon infection with virulent *P. syringae* (*Pst DC3000*) (Ward et al., 2010). Additionally, during ETI against *Psm* ES4326/*AvrRpt*2 (*P. syringe* pv *maculicola* strain ES4326, carrying the AvrRpt2 effector)—phenylalanine, asparagine, glutamine, and isoleucine significantly increase, while alanine, glutamate, leucine, tyrosine, and serine decrease (Yoo et al., 2020), suggesting that different amino acids contribute differently to plant immune responses.

Beyond their endogenous fluctuations, amino acids have been shown to trigger immune responses when applied exogenously. For instance, proline and threonine treatments induce hypersensitive-response-like cell death, restricting pathogen spread (Deuschle et al., 2004; Stuttmann et al., 2011). Similarly, glutamate application enhances resistance to rice blast disease by activating the salicylic acid (SA) pathway (Kadotani et al., 2016), and phenylalanine application induces PCD during ETI against *Psm* ES4326/*AvrRpt*2 (Yoo et al., 2020). These findings show that amino acids are active participants in plant immunity, yet the molecular mechanisms underlying their specific contributions remain largely unknown.

In this study, we identify serine as a novel defense activator. Serine pretreatment significantly delays PCD during ETI, suggesting a direct role in immune signaling. Serine is perceived by glutamate receptor-like calcium channel 3.4 (GLR3.4), which subsequently activates calcium-dependent protein kinase 5 (CPK5). Once activated, CPK5 phosphorylates transcription factors, including WRKY33, leading to the transcriptional activation of key defense genes. Consistent with this pathway, the serine-induced delay in PCD is largely compromised in *glr3.4* and *cpk5cpk6* mutants, confirming the essential role of the GLR3.4-CPK5-WRKY33 signaling cascade in serine-mediated defense priming. Our findings provide new insights into how primary metabolites contribute to plant immunity, expanding our understanding of the regulatory roles of amino acids beyond their traditional metabolic functions. Additionally, these findings suggest that amino acids can serve as natural defense activators, with potential applications in improving crop resistance to pathogens without the need for genetic engineering.

## Results

### Serine spray significantly delays PCD

To identify amino acids that influence PCD during ETI, we screened 18 amino acids, excluding tyrosine and tryptophan due to their low water solubility. To ensure that exogenous amino acids would not directly impact pathogen activity, we used transgenic Arabidopsis lines expressing a dexamethasone (DEX)-inducible *AvrRpt2* transgene in both the wild-type (WT) and *rps2* (AvrRpt2 receptor mutant) backgrounds, referred to as the WT-DEX *AvrRpt2* and *rps2*-DEX *AvrRpt2* lines (Mindrinos et al., 1994). Each amino acid was applied at a concentration of 5 mM, in accordance with previous reports on phenylalanine, proline, and threonine (Yoo et al., 2020; Deuschle et al., 2004; Stuttmann et al., 2011). We focused on amino acids that consistently yielded notable differences in PCD in pretreated plants versus water-sprayed control plants. Among all amino acids tested, serine resulted in the most significant PCD delays when applied 24 hours before DEX treatment (**Figure 1A, upper panel, and 1B**). Notably, serine had no effect in the *rps2* background, demonstrating that its impact on PCD depends on AvrRpt2 recognition (**Figure 1B**). The delayed PCD phenotype was visually apparent, which is particularly striking given that visible phenotypes are typically subtle 24 hours after DEX treatment (**Figure 1C**).

**Figure 1.**
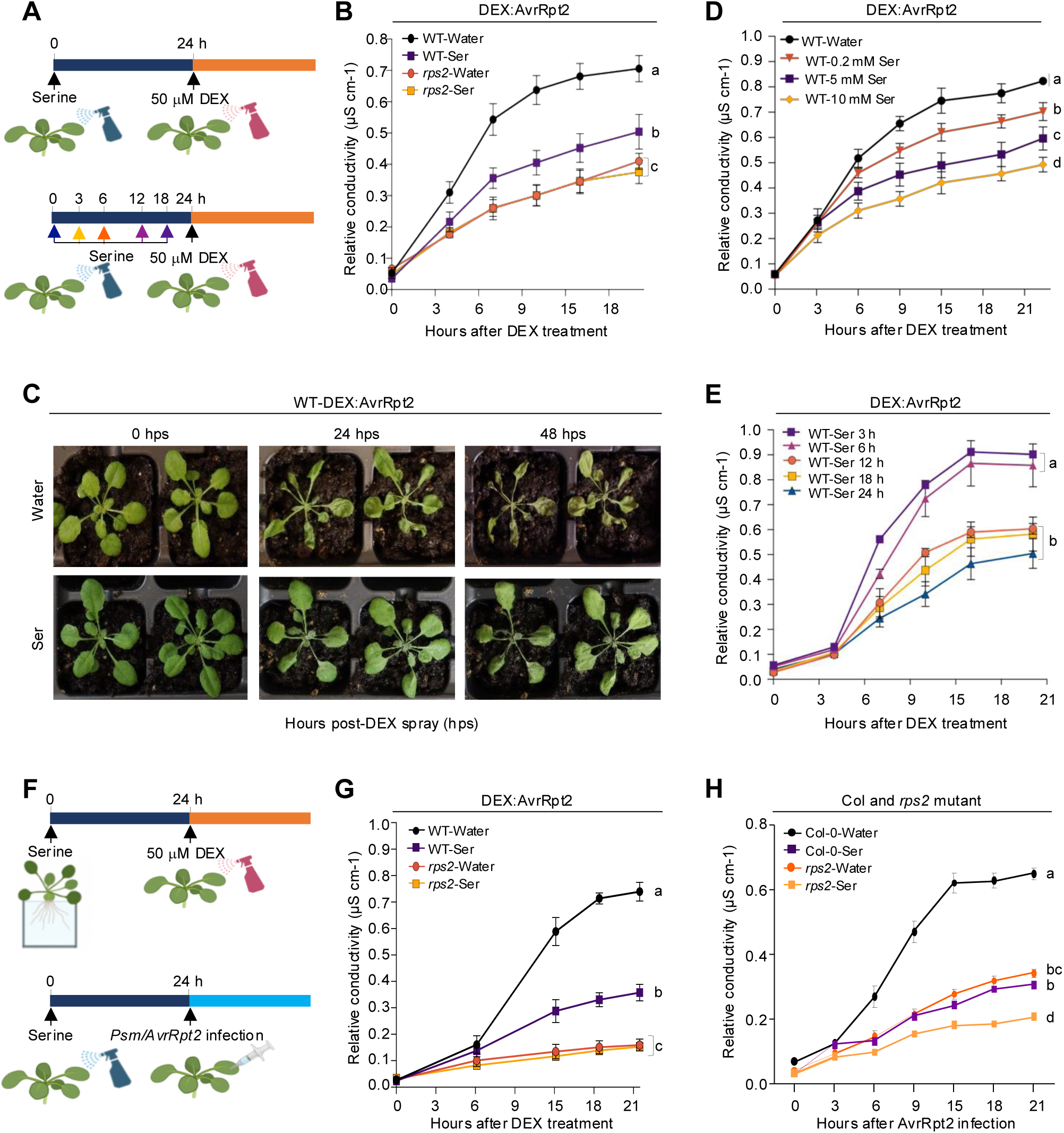
Exogenous serine application significantly delays PCD in a time- and concentration-dependent manner. **(A)** Schematic representation of the serine and DEX treatments used for the ion leakage experiments in panels **B**, **D**, and **E**. Water was used instead of serine for the control experiments. *Top panel*: For the results in panels **B** and **D**, DEX:AvrRpt2 plants were sprayed with water or serine, followed by a 50 μM DEX spray 24 hours later to trigger PCD. *Bottom panel*: For the results in panel **E**, plants were treated with serine, and then, after 3, 6, 12, 18, or 24 hours, DEX was applied to examine the time-interval-dependent effect of serine on the PCD delay. **(B)** Ion leakage analysis showing the effect of serine treatment. Conductivity was measured at the time points indicated on the x-axis. *n* = 4. **(C)** Representative images showing the PCD phenotype in WT-DEX:AvrRpt2 plants at 0, 24, and 48 hours post-DEX spray (hps). **(D)** Ion leakage analysis assessing the concentration-dependent effect of serine on PCD. Conductivity was measured at the time points indicated on the x-axis. *n* = 4. **(E)** Ion leakage analysis showing the effect of different time intervals between serine spray and DEX treatment. Conductivity was measured at the time points indicated on the x-axis. *n* = 3. **(F)** Schematic representation of the experimental conditions used to test the effects of serine by root-feeding (top panel) and to test the effectiveness of serine treatment against pathogen infection (bottom panel). *Top panel*: For the results in panel **G**, DEX:AvrRpt2 plants were root-fed with water or 5 mM serine, followed by 50 μM DEX spray 24 hours later. *Bottom panel*: For the results in panel **H**, leaves were sprayed with water or 5 mM serine, then infected with *Psm*/*AvrRpt2* (OD = 0.02) 24 hours after treatment. **(G)** Ion leakage analysis of plants in the root-fed serine assay. Conductivity was measured at the time points indicated on the x-axis. *n* = 2. **(H)** Ion leakage analysis showing the effect of exogenous serine treatment on WT and *rps2* plants. Conductivity was measured at the time points indicated on the x-axis. *n* = 3. For the ion leakage assays in (**B, D, E, G**, and **H)**, ion leakage values were normalized by dividing each measurement by the conductivity obtained after boiling the leaf discs, eliminating the variation in maximum conductivity. The data in (**B, D, E, G,** and **H**) represent the mean ± SEM of the number of biological replicates indicated for each panel. Statistical significance was determined using one-way ANOVA followed by Tukey’s *post hoc* test, for the measurements at the final time point.

To further explore this effect, we tested various serine concentrations (0.2, 5, and 10 mM) and observed a concentration-dependent delay in PCD. The weakest effect was detected at 0.2 mM, and the most pronounced delay occurred at 10 mM (**Figure 1D**). Next, we investigated the kinetics of the serine-induced PCD delay by varying the interval between serine application and DEX treatment (3−24 hours) (**Figure 1A, lower panel**). A 12-hour interval was required for a detectable effect, with the strongest PCD delay observed at 24 hours (**Figure 1E**). These findings indicate that serine modulates PCD in a time- and concentration-dependent manner.

To determine whether serine acts locally or systemically, we compared foliar spraying and root feeding (**Figure 1A and 1F**, upper panels). Root-fed serine delayed PCD to a similar extent as leaf-sprayed serine (**Figure 1B and 1G**), suggesting that serine or a serine-activated signal is transported systemically, likely via the vasculature, to regulate PCD regardless of the application method.

Finally, we tested whether exogenous serine could also delay PCD during pathogen infection. Wild-type Col-0 plants were pretreated with 5 mM serine before *Psm*/*AvrRpt2* infection, revealing a significant PCD delay (**Figure 1H**). Interestingly, a moderate PCD delay was also observed in the *rps2* mutant, despite its lack of ability to recognize AvrRpt2. Since *rps2* mutants exhibit residual PCD during pathogen infection–unlike *rps2*-DEX *AvrRpt2* plants, which show no PCD–the observed delay in the *rps2* mutants may indicate the existence of alternative mechanisms of PCD that function independently of AvrRpt2 recognition. Collectively, these findings demonstrate that serine pretreatment delays PCD in a concentration- and time-dependent manner, acting both locally and systemically.

### Serine treatment primes defense gene expression

Given the observed serine-induced PCD delay, we investigated whether serine could activate defense gene expression in the absence of pathogen infection and whether this activation contributes to the PCD delay. Using wild-type Col-0 plants, we analyzed the expression of key defense marker genes at multiple time points (0, 0.5, 1, 6, and 24 hours) following the foliar application of 5 mM serine (**Figure 2A and 2B**). Notably, all tested defense-related genes, including *NDR1*/*HIN1-LIKE 10* (*NHL10*)*, PHYTOALEXIN DEFICIENT 3* (*PAD3*), *ISOCHORISMATE SYNTHASE 1* (*ICS1*), *PATHOGENESIS*-*RELATED 1* (*PR1*), *AGD2*-*LIKE DEFENSE RESPONSE PROTEIN 1* (*ALD1*), and *FLAVIN*-*DEPENDENT MONOOXYGENASE 1* (*FMO1*) (Zhou et al., 1999; Dubiella et al., 2013; Yang et al., 2015; Holmes et al., 2019), exhibited significant upregulation 24 hours post-serine application. However, no significant changes were detected at earlier time points (0.5, 1, or 6 hours) or in water-treated controls (**Figure 2B**). These results suggest that serine functions as a defense activator, but more than 6 hours is required for it to induce defense gene expression. Consistent with these observations, serine treatment led to significant SA accumulation 24 hours post-application, correlating with the induction of *ICS1*, a key gene in SA biosynthesis (**Figure 2C**).

**Figure 2.**
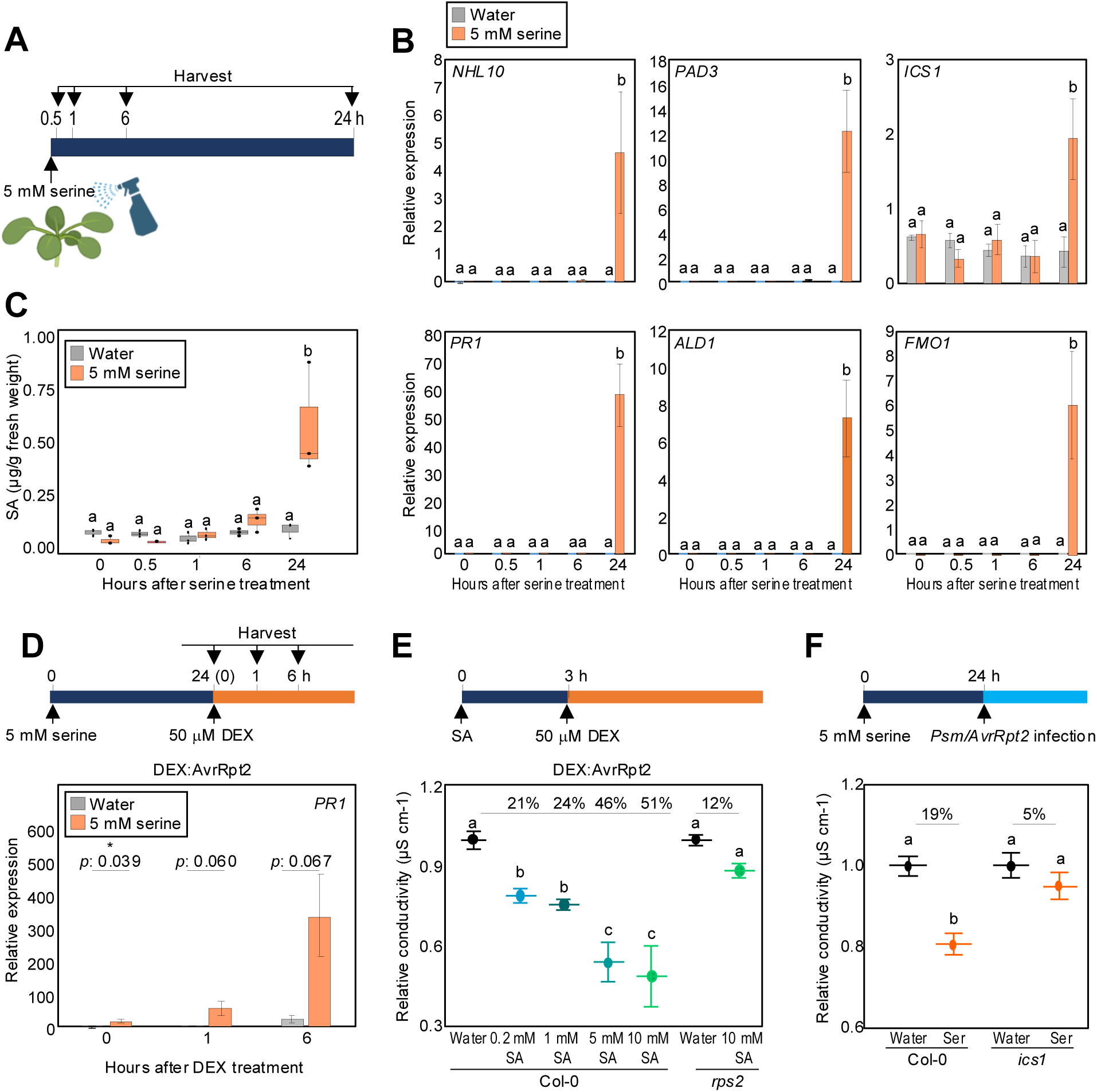
Serine spray induces defense genes and delays PCD. **(A)** Schematic representation of the time points for sample collection following 5 mM serine spray (0, 0.5, 1, 6, and 24 hours). **(B)** RT-qPCR analysis of key defense gene expression after serine treatment. Leaves were sprayed with water or 5 mM serine and harvested at the indicated time points. *n* = 3. **(C)** Quantification of SA levels following serine treatment. *n* = 3. **(D)** *PR1* expression in WT-DEX::AvrRpt2 plants pretreated with serine or water, followed by 50 μM DEX spray 0, 1, or 6 hours later. *n* = 3. **(E)** Ion leakage analysis of DEX::AvrRpt2 plants pretreated with water or various concentrations of SA (0.2 mM, 1 mM, 5 mM, and 10 mM). Leaves were sprayed with water or SA, followed by a 50 μM DEX spray 3 hours later. Conductivity was measured 1 day after the DEX spray. Percent values indicate the conductivity relative to that of water-treated plants. *n* = 3. **(F)** Ion leakage analysis comparing WT and *ics1* mutant plants pretreated with 5 mM serine or water, then infected with *Psm*/*AvrRpt2* (OD = 0.02) 24 hours after the pretreatment. *n* = 3. Conductivity was measured 1 day after the DEX spray. Percent values indicate the conductivity relative to that of water-treated plants. For the ion leakage assays in **(E)** and **(F),** ion leakage values were normalized by dividing each measurement by the conductivity obtained after boiling the leaf discs, eliminating the variation in maximum conductivity. The data in **(B, D, E,** and **F)** represent the mean ± SEM of the number of biological replicates indicated for each panel. The data in **(C)** are presented as box and whisker plots. Statistical significance for (**B**, **C**, **E**, and **F**) was determined using one-way ANOVA followed by Newman–Keuls’ *post hoc* test. Different letters indicate significant differences (*p* < 0.05). Statistical significance for **(D)** was assessed using Student’s *t*-test. The asterisk indicates a statistically significant difference (\**p* < 0.05).

Given the simultaneous increase in defense gene induction and SA accumulation, we hypothesized that serine pretreatment primes the plant’s defense responses, enabling quicker and stronger defense activation upon pathogen infection. To test this hypothesis, we pretreated WT-DEX *AvrRpt2* plants with 5 mM serine or water for 24 hours, followed by DEX treatment to induce *AvrRpt2* expression. Prior to DEX treatment, *PR1* expression was already significantly higher in serine-pretreated plants than in water-pretreated controls (**Figure 2D**), consistent with our findings in wild-type Col-0 plants (**Figure 2B**). One hour after DEX treatment, *PR1* expression further increased in serine-pretreated plants, whereas no significant induction was observed in water-pretreated plants. By 6 hours post-DEX spray, *PR1* expression had increased after both pretreatments but remained significantly higher in serine-pretreated plants (**Figure 2D**). These results confirm that serine pretreatment primes defense responses, enabling a more rapid and robust activation of defense genes upon pathogen effector recognition.

To determine whether defense gene priming mediates serine-induced PCD suppression, we induced the priming effect using SA pretreatment. To test whether exogenous SA could mimic serine’s effect within a shorter timeframe, plants were pretreated with SA for 3 hours before DEX application to induce AvrRpt2-triggered PCD. SA pretreatment significantly delayed PCD, mirroring the effect of serine (**Figure 2E**). To further validate this connection, we tested serine pretreatment in the *ics1* mutant, which is deficient in SA biosynthesis. Unlike wild-type plants, the *ics1* mutant failed to exhibit a PCD delay following serine pretreatment (**Figure 2F**), confirming that SA-dependent defense gene priming is essential for the serine-induced PCD delay. Collectively, these findings demonstrate that serine functions as a defense priming signal, activating SA-dependent defense gene expression and enhancing immune priming. This priming effect enables plants to mount a faster and more-robust defense response upon pathogen attack, ultimately delaying PCD.

### GLR3.4 contributes to the serine-dependent activation of defense-responsive genes

To investigate how exogenous serine activates defense gene expression, and to identify amino acid receptors involved in serine perception, we examined the role of glutamate-like receptor (GLR) proteins, which function as amino-acid-gated ion channels. Given that serine-induced changes in membrane potential are disrupted in *glr3.3* and *glr3.4* mutants (Stephens et al., 2008; Forde and Roberts, 2014; Green et al., 2021), we hypothesized that GLR receptors mediate serine-induced defense activation. To test this possibility, we analyzed defense gene expression in wild-type, *glr3.3*, and *glr3.4* plants following 5 mM serine treatment. While *glr3.3* mutants showed no significant difference in defense gene induction compared to wild-type plants, *glr3.4* mutants exhibited a marked reduction in serine-induced defense gene expression (**Figure 3A**). These results indicate that GLR3.4, but not GRL3.3, is required for the serine-mediated activation of defense pathways. To determine whether GLR3.4 is also necessary for the serine-induced PCD delay, we pretreated wild-type Col-0, *glr3.3*, and *glr3.4* plants with 5 mM serine or water, then infected them with *Psm*/*AvrRpt2*. Serine significantly delayed PCD in the wild type and in *glr3.3* mutants, but this effect was partially compromised in *glr3.4* mutants (**Figure 3B**). These findings demonstrate that GLR3.4 is the primary receptor for serine, triggering downstream defense activation and contributing to the serine-mediated delay in PCD.

**Figure 3.**
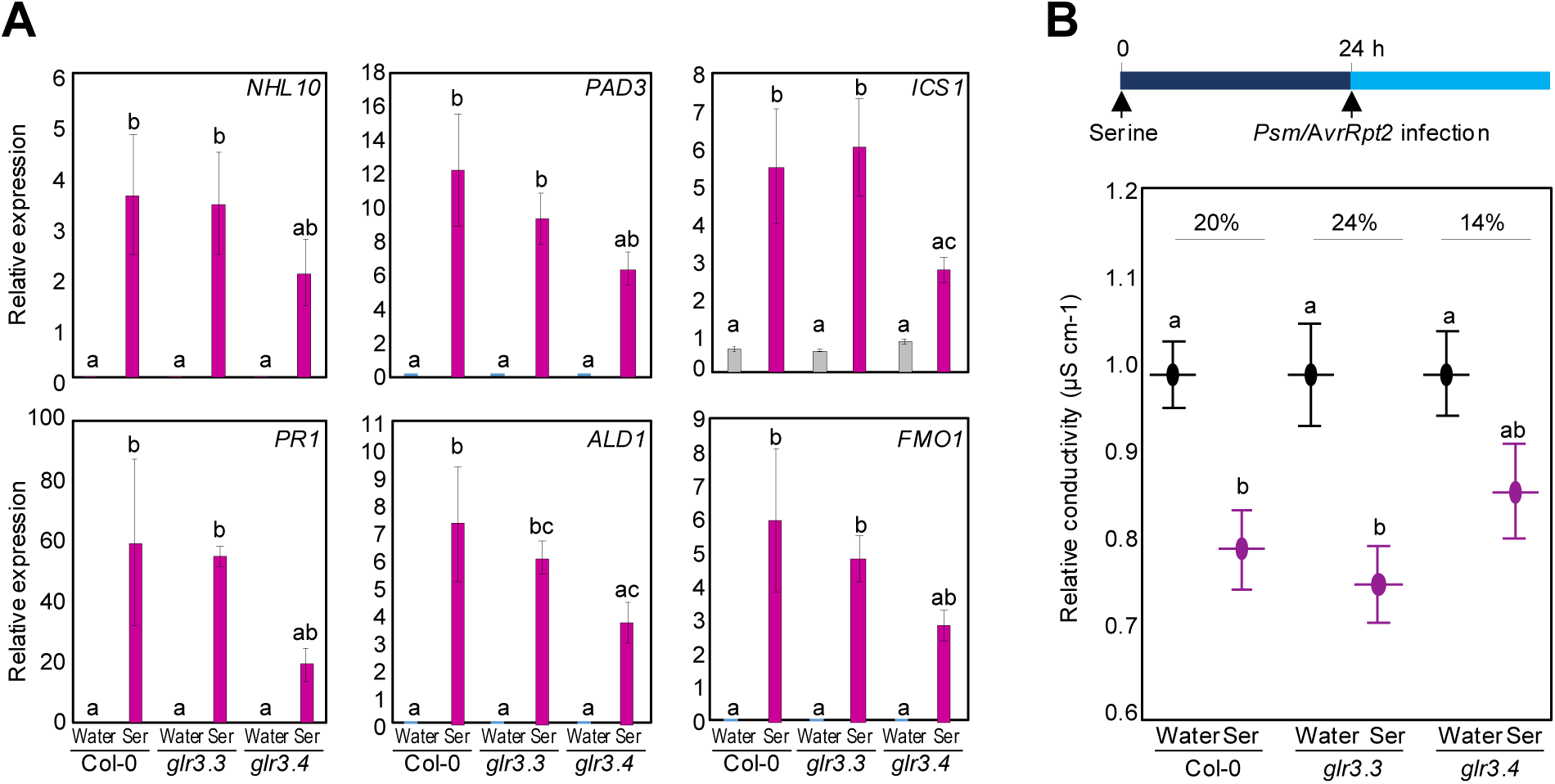
GLR3.4 contributes to the serine-dependent activation of stress-responsive genes. **(A)** RT-qPCR analysis of key defense genes in wild-type Col-0, *glr3.3*, and *glr3.4* mutants following 5 mM serine treatment. *n* = 3. **(B)** Ion leakage analysis of wild-type Col-0, *glr3.3*, and *glr3.4*. Leaves were sprayed with water or 5 mM serine, then infected with *Psm*/*AvrRpt2* (OD = 0.02) 24 hours after pretreatment. Conductivity was measured 1 day after the DEX spray. The results from the water treatment set to 1 and the results from the serine treatment normalized relative to water for each genotype. Percent values indicate the difference relative to water-treated plants. *n* = 8. The data in **(A and B)** represent the mean ± SEM of the number of biological replicates indicated for each panel. Statistical significance was determined using one-way analysis ANOVA followed by Newman–Keuls’ *post hoc* test. Different letters indicate significant differences (*p* < 0.05).

### CPK5 is required for the serine-dependent activation of defense genes via its kinase activity

To determine whether serine-induced defense gene activation depends on calcium signaling downstream of GLR3.4, we pretreated wild-type plants with EGTA, a calcium chelator (Ma et al., 2017; Shin et al., 2023), in the presence or absence of serine. Serine-induced defense gene expression was lower in the presence of EGTA, indicating that calcium signaling is essential for serine-mediated defense activation (**Figure 4A**). Given this calcium dependence, we examined CPKs as a potential downstream signaling component. Previous studies have identified CPK5 as a key regulator of defense gene activation in response to pathogen effectors (Dubiella et al., 2013; Gao et al., 2013; Gao and He, 2013). To determine whether CPK5 contributes to serine-induced defense activation, we examined serine-induced defense gene expression in wild-type plants and in the *cpk5cpk6* double mutant, as CPK5 and CPK6 have partially overlapping functions (Zhou et al., 2020). Most defense genes, except *ICS1*, showed significantly less induction in *cpk5cpk6* mutants than in wild-type plants following serine treatment (**Figure 4B**). Consequently, the serine-induced delay of PCD was partially compromised in *cpk5cpk6* mutants (**Figure 4C**), further supporting the role of CPK5 and CPK6 in serine-mediated defense activation. To further dissect the role of CPK5’s kinase activity in serine-mediated defense gene activation, we used a *DEX*:*CPK5ac-dead* line, which carries a D221A point mutation in the CPK kinase domain, rendering it catalytically inactive (Yip Delormel et al., 2022). DEX treatment in this line induces the overexpression of kinase-dead *CPK5*, allowing us to specifically assess whether its kinase activity is necessary for serine signaling. Unlike *cpk5cpk6* mutants, where *ICS1* expression remained unchanged (**Figure 4C**)–likely due to compensation by other CPKs (Boudsocq et al., 2010)–the serine-induced activation of *ICS1* and other defense genes was significantly reduced in the kinase-dead line (**Figure 4D**). This result suggests that inactive CPK5 may interfere with the functions of other CPKs, preventing *ICS1* induction. In contrast, plants without DEX treatment, which retained functional CPK5, exhibited robust defense gene induction (**Figure 4D**). To further confirm the role of CPK5 in serine-mediated PCD regulation, we examined the PCD phenotype in the kinase-dead CPK5 line following serine treatment. In the absence of DEX, serine delayed PCD, similar to the response observed in wild-type Col-0 plants. However, in the presence of DEX, the serine-induced PCD delay was compromised (**Figure 4E**), confirming that CPK5 kinase activity is essential for serine-mediated immune priming. Collectively, these results demonstrate that CPK5 functions as a critical signal transducer in serine-mediated defense responses.

**Figure 4.**
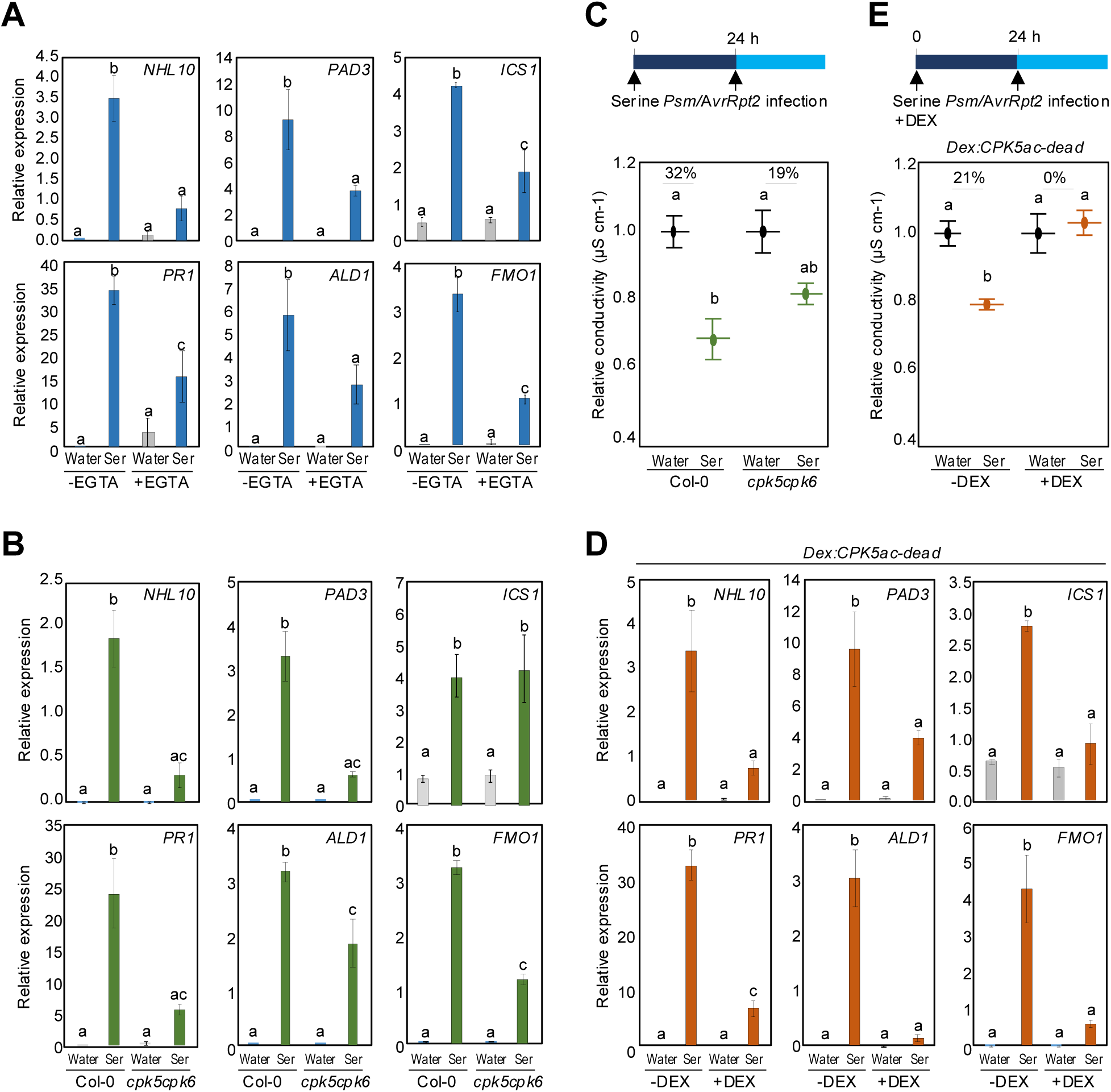
CPK5 is essential for serine-mediated defense gene activation. **(A)** RT-qPCR analysis of key defense genes in wild-type Col-0 treated with 5mM serine alone or 5 mM serine + 0.5 mM EGTA. *n* = 3. **(B)** RT-qPCR analysis of key defense genes in *cpk5cpk6* plants following 5 mM serine treatment. *n* = 3. **(C)** Ion leakage analysis of wild-type Col-0 and *cpk5cpk6* plants. Leaves were sprayed with water or 5 mM serine, then infected with *Psm*/*AvrRpt2* (OD = 0.02) 24 hours after pretreatment. Conductivity was measured 1 day after the DEX spray. The results from the water treatment set to 1 and the results from the serine treatment normalized relative to water for each genotype. Percent values indicate the difference relative to water-treated plants. *n* = 3. **(D)** RT-qPCR analysis of key defense genes in *DEX:CPK5-ac-dead* plants following 5 mM serine treatment with or without DEX. *n* = 3. **(E)** Ion leakage analysis in *DEX:CPK5-ac-dead* plant following serine pretreatment with or without DEX. Leaves were sprayed with water or 5 mM serine, with or without 50 μM DEX, then infected with *Psm*/*AvrRpt2* (OD = 0.02) 24 hours later. Conductivity was measured 1 day after the DEX spray. The results from the water treatment set to 1 and the results from the serine treatment normalized relative to water for each genotype. Percent values indicate the difference relative to water-treated plants. *n* = 3. Data in **(A-E)** represent the mean ± SEM of the number of biological replicates indicated for each panel. Statistical significance was determined using one-way ANOVA followed by Newman–Keuls’ *post hoc* test. Different letters indicate significant differences (*P* < 0.05).

### WRKY33 mediates the serine-dependent activation of defense genes downstream of CPK5

Previous studies have demonstrated that CPK5 phosphorylates several WRKY transcription factors, including WRKY33, a known regulator of plant immunity (Gao et al., 2013; Zhou et al., 2020;Yamada & Mine, 2024). To assess the role of WRKY33 in serine-induced defense signaling downstream of CPK5, we analyzed defense gene expression in the *wrky33* mutant following serine treatment. The mutant showed significantly reduced induction of *NHL10, PAD3, ALD1*, and *FMO1*, while the expression of *ICS1* and *PR1* remained unchanged (**Figure 5A**). These results further suggest that the CPK5-WRKY33 signaling module plays a critical role in mediating the serine-induced activation of specific defense genes.

**Figure 5.**
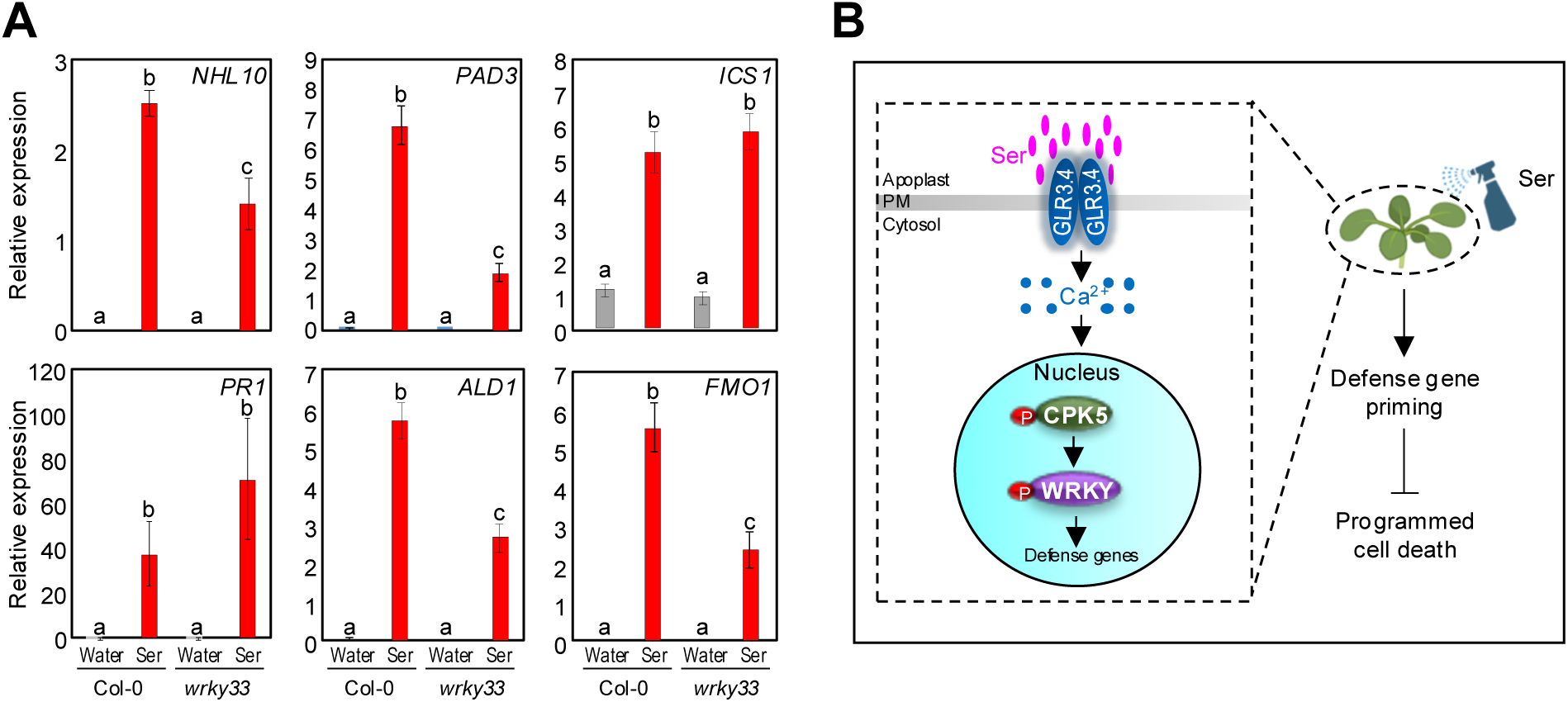
WRKY33 mediates serine-induced defense gene activation and proposed model for GLR3.4–CPK5–WRKY signaling in immune priming. **(A)** RT-qPCR analysis of key defense genes in wild-type Col-0 and *wrky33* mutants following 5 mM serine treatment. Data represent the mean ± SEM of 3 biological replicates. Statistical significance was determined using one-way ANOVA followed by Newman–Keuls’ *post hoc* test. Different letters indicate significant differences (*p* < 0.05). **(B)** Proposed model for serine-induced defense activation. Pretreatment with serine primes the plant by activating defense gene expression prior to pathogen attack. Serine activates GLR3.4, triggering a calcium influx and activating CPK5, which phosphorylates WRKY33. Activated WRKY33, potentially along with other WRKY transcription factors or additional CPK5 targets, upregulates defense genes, enhancing immune readiness and leading to a stronger and faster response upon pathogen attack. This primed state helps delay pathogen-induced PCD, improving protection during infection.

Taken together, our findings establish a mechanistic framework for how exogenous serine regulates plant defense responses and delays PCD during ETI. Upon serine treatment, GLR3.4, an ionotropic glutamate receptor-like channel on the plasma membrane, perceives serine and triggers a calcium-dependent signaling cascade. This signaling cascade leads to the activation of CPK5. Activated CPK5 subsequently phosphorylates WRKY33, which acts as a critical downstream component. WRKY33, in turn, activates the transcription of specific defense-related genes, resulting in enhanced immune responses and delayed PCD upon pathogen attack (**Figure 5B**).

## Discussion

Plants rely on intricate regulatory networks to balance growth and defense while responding to pathogen attack. Certain chemical inducers can activate endogenous defense mechanisms, mimicking immune signals and priming immunity. For example, non-metabolizable glucose (Glc) analogs, such as 3-O-methyl-d-Glc and 2-deoxy-d-Glc (2DG), enhance defense gene expression by promoting CPK5 phosphorylation (Yamada and Mine, 2024). Similarly, SA analogs, such as 2,6-dichloroisonicotinic acid (INA) and benzothiadiazole (BTH), trigger SA-responsive gene expression (Ward et al., 1991; Uknes et al., 1992; Palmer et al., 2019). Additionally, 3,5-dichloroanthranilic acid (DCA) triggers NONEXPRESSOR OF PATHOGENESIS-RELATED GENES 1 (NPR1)-dependent immune signaling (Knoth et al., 2009). Understanding the regulatory roles of such chemical inducers provides valuable insights into how immune signaling pathways are modulated, with potential applications for improving disease resistance in plants.

Amino acids are increasingly recognized as important regulators of plant immunity. For example, proline application induces a calcium-mediated oxidative burst and enhances SA synthesis, while glutamate stimulates pathogen-associated molecular pattern (PAMP)-induced gene expression (Chen et al., 2011; Goto et al., 2020). However, the precise mechanisms through which specific amino acids act as defense activators remain poorly understood. Our study identifies serine as a novel immune regulator that functions via the GLR3.4-CPK5-WRKY33 signaling pathway. Serine-induced defense activation was significantly impaired in *glt3.4, cpk5cpk6,* and *wrky* mutants, demonstrating the requirement for this signaling cascade. GLRs are increasingly recognized as amino acid sensors in plants: GLR3.1 and GLR3.5 respond to methionine (Kong et al., 2016; Ju et al., 2020), while GLR3.3 binds to glutamate, cysteine, and methionine (Alfieri et al., 2020; Grenzi et al., 2021). Previous studies have demonstrated that GLR3.4 is required for serine-induced changes in membrane potential (Stephens et al., 2008). Consistent with this observation, our findings establish GLR3.4 as necessary for serine-mediated immune activation and delayed PCD. However, since serine-induced defense activation was not entirely abolished in *glr3.4* mutants, it is likely that additional GLR receptors or alternative signaling components contribute to serine perception. Given the functional redundancy among GLR receptors in other physiological processes (Nguyen et al., 2018; Xue et al., 2022), future studies should investigate which GLR combinations or other signaling components function alongside GLR3.4 in serine-mediated immune responses.

CPK5 is a well-characterized calcium-dependent kinase that plays a pivotal role in ETI (Gao et al., 2013). During ETI, pathogen effector recognition triggers a cytosolic calcium influx, leading to CPK5 activation. Once activated, CPK5 translocates to the nucleus, where it phosphorylates WRKY transcription factors, enhancing their DNA binding activity (Gao et al., 2013). A recent study demonstrated that CPK5 is required for sucrose-mediated defense gene activation, as the glucose-6-phosphate (G6P)-mediated suppression of protein phosphatases promotes CPK5 phosphorylation and primes immune responses before pathogen infection (Yamada and Mine, 2024). Consistent with this regulatory role, our study shows that CPK5 mediates serine-induced defense gene activation, reinforcing its central function in plant immune signaling. Furthermore, our study confirms that WRKY33, a known CPK5 target (Zhou et al., 2020), is essential for serine-induced defense gene activation. Interestingly, *ICS1* and *PR1* expression remained unchanged in *wrky33* mutants. This observation suggests that in addition to WRKY33, other transcription factors or regulatory proteins downstream of CPK5 may mediate the serine-induced activation of *ICS1* and *PR1*, warranting further investigation.

PCD is a hallmark of ETI, which is triggered when nucleotide-binding leucine-rich repeat (NB-LRR) receptors detect pathogen effectors (Coll et al., 2011). Our findings demonstrate that serine pretreatment delays PCD by priming defense gene expression. Similarly, SA pretreatment delays PCD, and notably, the serine-induced PCD delay is abolished in the *ics* mutant, which is deficient in SA biosynthesis. A previous study showed that defense priming via flg22 treatment restricts the accumulation of the type III effector AvrPto, with significantly reduced *AvrPto* levels in flg22-treated leaves infected with *PstDC3000* (Rogan et al., 2024). These results suggest that serine-mediated defense priming may restrict AvrRpt2 accumulation, thereby delaying RPS2-dependent recognition and PCD onset.

Since serine serves as a precursor for multiple metabolites, it is possible that serine-derived metabolites contribute to immune responses, either independently or in conjunction with serine itself. While our findings establish serine as a key metabolite responsible for defense gene activation, its downstream metabolites may function at different time points or through distinct regulatory mechanisms. Future studies utilizing labeled serine will be essential for tracking its metabolic fate and determining whether its downstream metabolites directly regulate plant immunity or contribute indirectly through secondary signaling pathways.

Defense activators are promising tools for enhancing crop resilience against pathogens (Goto et al., 2020; Marolleau et al., 2017). Unlike conventional pesticides, which pose risks such as the emergence of resistant pathogen strains and environmental toxicity, chemical inducers that activate plant immunity offer a safer and more sustainable alternative (Goto et al., 2020; Marolleau et al., 2017). Our findings demonstrate that exogenously applied serine can function as immune priming agent, supporting its potential use in agriculture as a foliar spray to enhance disease resistance without the need for genetic modification. Future research should focus on optimizing serine application protocols to maximize its efficacy across diverse crop species and environmental conditions. Additionally, investigating the broader role of amino-acid-mediated plant defense regulation could provide deeper insights into how plants balance growth and defense.

## Materials and methods

### Plant materials and growth conditions

*Arabidopsis thaliana* plants were grown in Sunshine Complete Mix #1 soil (Sun Gro Horticulture, Agawam, MA, USA) in a walk-in growth room under controlled conditions (22 °C, 55% humidity, and a 12-h/12-h light/dark cycle). The *ics1* (SALK_042603), *glr3.3* (SALK_099757), *glr3.4* (SALK_079842), *cpk5cpk6* (CS69905), and *wrky33* (SALK_006603) seeds were obtained from the Arabidopsis Biological Resource Center (ABRC) previously characterized in published studies (Zheng et al., 2006; Gao et al., 2013; Sasek et al., 2014; Salvador-Recatala, 2016; Cheng et al., 2018). and *DEX:CPK5-ac-dead* seeds were generously provided by Dr. Marie Boudsocq and previously described in Yip Delormel et al., 2022.

### Exogenous application of serine, SA, DEX, and EGTA

For serine spray treatments, 0.2, 5, and 10 mM serine solutions were prepared by dissolving serine (Sigma-Aldrich, S4500) in autoclaved Milli-Q water. Autoclaved Milli-Q water was used as a control. For SA spray treatments, 0.2, 1, 2, 5, and 10 mM SA solutions were prepared by dissolving SA (Sigma-Aldrich,71945) in autoclaved Milli-Q water. For DEX spray treatments, a 500X stock solution was prepared by dissolving DEX powder (Sigma-Aldrich, D4902) in 10 mL of 100% ethanol. This stock was then diluted to a 1X working solution in autoclaved Milli-Q water to a final concentration of 50 μM before application. For EGTA spray treatments, a 0.5 mM EGTA solution was prepared by dissolving EGTA (Sigma-Aldrich, 324626) in autoclaved Milli-Q water. Leaves of 4-week-old plants were sprayed with each solution. For the root-feeding experiments, 4-week-old plants were carefully removed from the soil and transferred to containers containing either water or 5 mM serine solution. One day later, plants were replanted in soil before DEX treatment.

### Ion leakage measurements

PCD was quantified using ion leakage assays as described previously (Hatsugai and Katagiri, 2018). For DEX-inducible AvrRpt2 transgenic lines (WT-DEX *AvrRpt*2 and *rps2*-DEX *AvrRpt2*), plants were pretreated with serine or SA spray, followed by treatment with 50 μM DEX to induce AvrRpt2 expression and PCD activation. For wild-type and mutant plants, leaves were infiltrated with *Psm* ES4326/*AvrRpt2* (OD_600nm_ = 0.02) instead of DEX treatment. One hour post-DEX treatment or *Psm* ES4326/*AvrRpt2* infection, leaf discs (5 per sample) were washed and placed in 5 mL of fresh water in a 50 mL Corning tube. Electrolyte leakage was measured at 3-4-hour intervals over 24 hours using a conductivity meter (VWR, STAR A322).

### RNA extraction and quantitative real-time polymerase chain reaction (qRT-PCR)

Total RNA was extracted using a Quick-RNA Miniprep Kit (Zymo Research). cDNA was synthesized from 1 μg total RNA using LunaScript RT Supermix (New England BioLabs) following the manufacturer’s protocol. qRT-PCR was performed using Luna Universal qPCR Master Mix (New England BioLabs) on a LightCycler 480 real-time PCR system (Roche). Gene expression was normalized to *PP2A* (At1g13320) expression (Klie and Debener, 2011), and relative expression levels were calculated using the comparative ΔΔCt method. Primers are listed in Supplementary Data Set S1.

### SA measurement

SA was extracted following a previously described method (Zheng et al., 2015). Approximately 200 mg of leaf tissue was extracted using 90% (vol/vol) methanol, followed by 100% methanol. Samples were vacuum-dried and resuspended in 5% (vol/vol) trichloroacetic acid. SA was extracted twice using a 1:1 ethyl acetate/cyclopentane mixture. The extracts were dried and dissolved in HPLC eluent (10% methanol in 0.2 M acetate buffer). SA levels were analyzed using an Agilent 1260 high-performance liquid chromatography system equipped with an XTerra MS C18 column (Waters, 3.5 µm, 4×100 mm).

## Statistical analysis

All statistical analyses were performed using GraphPad Prism 10 (GraphPad Software, La Jolla, CA, USA).

## Supporting information

Supplementary Data Set

## Acknowledgments

We thank Dr. Marie Boudsocq (Institute of Plant Sciences Paris Saclay, France) for providing *DEX:CPK5-ac-dead* seeds. We also thank the HYoo Lab members for their critical comments on this project. This study was supported by start-up funds from the University of Utah and Oklahoma State University.

## Author Contributions

K.L., A.N., and H.Y. designed the research. K.L. performed all experiments except for the following: A.N. conducted the prescreening of amino acids for PCD using ion leakage assays and performed the serine-concentration-dependent ion leakage analysis; J.C. conducted the time-dependent ion leakage assays and the root-fed serine experiments. All authors analyzed the data. K.L. and H.Y. wrote the manuscript with input from all authors.

